# Nature Forest Reserves in Tanzania and their importance for conservation

**DOI:** 10.1101/2023.01.24.525332

**Authors:** Claire Ract, Neil D. Burgess, Lars Dinesen, Peter Sumbi, Isaac Malugu, Julia Latham, Lucy Anderson, Roy E. Gereau, Marcelo Gonçalves de Lima, Amina Akida, Evarist Nashanda, Gertrude Lyatuu, Phil Platts, Francesco Rovero

## Abstract

Since 1997 Tanzania has undertaken a process to identify and declare a network of Nature Forest Reserves (NFRs) with high biodiversity values, mainly from within its existing portfolio of national Forest Reserves, but with some new extensions. In recent years this expansion has accelerated, with ten NFRs declared since 2015. The current network of 19 existing NFRs covered 918,212 hectares by the end of 2020 (with an additional three reserves covering 34,862 hectares in the process of being declared). The coverage by NFRs of Tanzanian habitat types has increased to include the main forest types, wet, seasonal, and dry, and includes wetlands and grasslands. This has led to more than a doubling of the coverage of species ranges of vertebrates by the NFR network. Declared and proposed reserves now contain at least 178 of Tanzania’s 242 endemic vertebrate species, of which over 50% are threatened with extinction, and 553 Tanzanian endemic plant taxa (species, subspecies, and varieties), of which 61.3% are threatened. NFRs also support 41 single site endemic vertebrate species and 76 single site endemic plant taxa. Time series data show that NFR management effectiveness is increasing, especially where donor funds are available. Tourism, diversified revenue generation and investment schemes are required to create a network of economically self-sustaining NFRs able to conserve critical biodiversity values. Improved management and investment have reduced some threats in recent years, but ongoing challenges include illegal logging, charcoal production, firewood, pole cutting, hunting, fire, wildlife trade, and the unpredictable impacts of climate change.

## Introduction

Protected areas are essential for conserving biodiversity on earth, and important for storing carbon and thereby an important buffer against climate change [1,2]. The International Union for Conservation of Nature (IUCN) defines a Protected Area (PA) as “An area of land and/or sea especially dedicated to the protection and maintenance of biological diversity, and of natural and associated cultural resources, and managed through legal or other effective means” [1].

The creation of protected forest areas in Tanzania has a long history stretching back to the German colonial period in the late 1800s. This ‘forest reserve’ network has expanded over time, first during the German and subsequent British colonial periods, and then since Tanzanian independence in 1961 [3, 4, 5, 6]. Management aims include reserves established for production from natural forests (timber and charcoal), protection of natural forests (water catchment reserves and for the prevention of landslides and erosion), and plantation forestry using exotic species.

After the implementation of the ‘new’ Forest Policy in 1998 [7] and the Forest Act in 2002 [8], forest reserves started to be designed for the preservation of their biodiversity and habitats, and human activities were restricted [4, 5]. Under this new legislative framework, Nature Forest Reserves (NFRs) were recognised as forest areas of particularly high importance for globally unique biodiversity and managed in most cases with strong protection. The first phase of declaring NFRs was in the Eastern Arc Mountains ecoregion [9] during the period 1997-2009, starting with Amani Nature Reserve in the East Usambara Mountains in 1997.

Over the past 25 years, the Tanzanian government has continued to work to identify and upgrade other biologically important reserves to become NFRs. These sites initially fell under the ownership and management of the Forestry and Beekeeping Division of the Ministry of Natural Resources and Tourism, and (since 2005) the Tanzania Forest Services (TFS) Agency. After the first phase of declaring NFRs, the network expanded to cover all the different forest types in the country (including the Coastal forests, the Northern Volcanic, the Southern Highlands, Guineo-Congolian, Miombo and Miombo-Acacia forests). The current network of NFRs in Tanzania contains 19 reserves, with an additional three reserves in the process of being declared (Figure 1).

**Fig 1:**
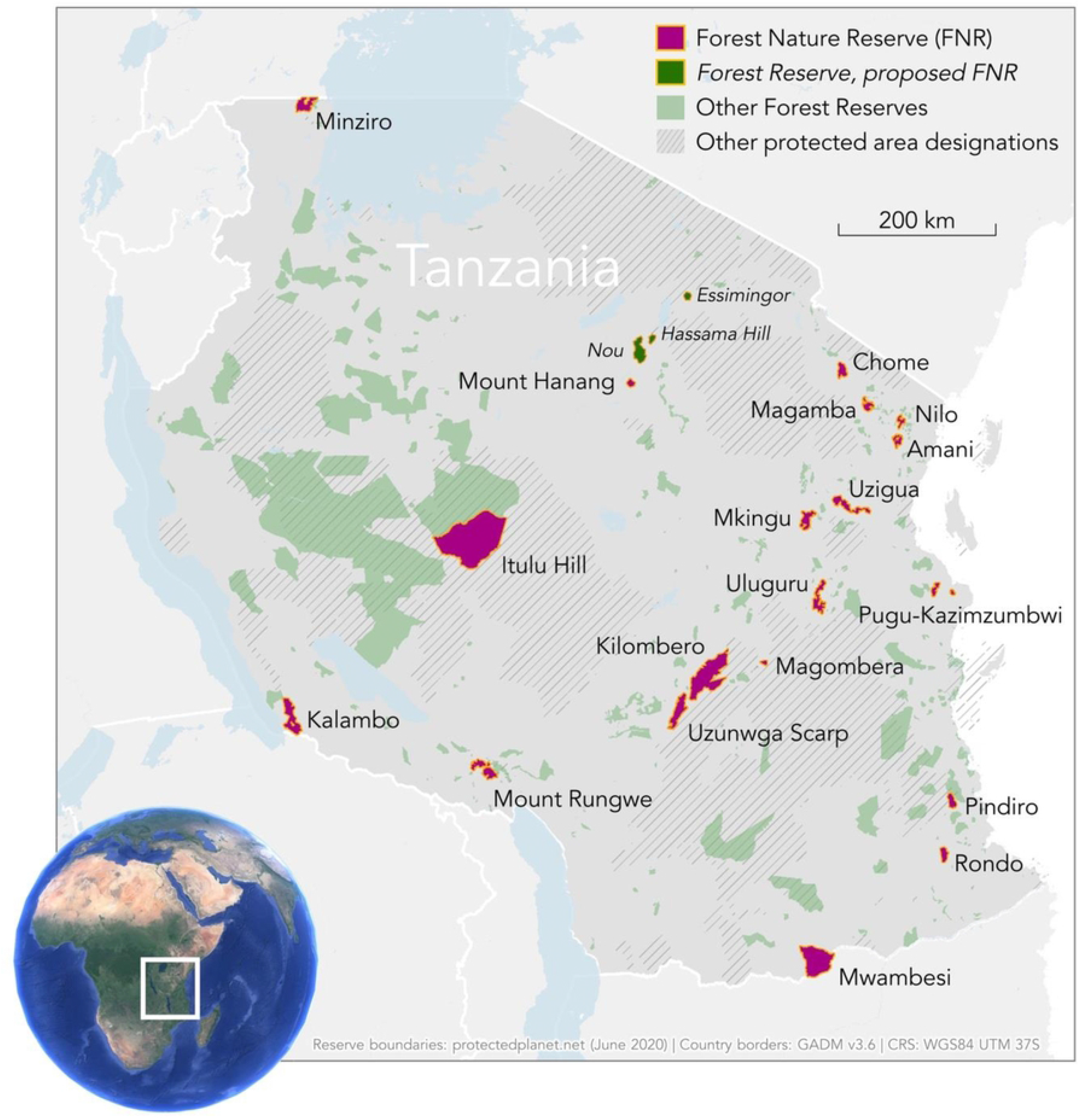
Map of Tanzania showing Nature Forest Reserves (NFRs, red), as well as Forest Reserves (green) including three in the process of being declared NFRs (italicized names), and other kinds of protected areas (shaded).

In this paper we describe Tanzania’s NFR network for the first time by: (i) evaluating the development of the NFR network in Tanzania and assessing its coverage of Tanzanian biodiversity values, (ii) assessing the effectiveness of NFRs and whether management resources have been deployed to the most important sites to achieve conservation goals, and (iii) assessing the coverage of endemic and rare species by the NFR network and other protected areas across Tanzania. We discuss the findings in light of our own insights into management challenges in the NFR and ways that these might be addressed in the future.

## Methods

### Data collection

Between September 2019 and November 2021, we collated available published and unpublished data on the Tanzanian NFRs (summarised in Table 1). Data on 19 NFRs declared and 3 proposed NFRs in or before 2020 included declaration details, biological features, management information, and threats. Management data on the five last NFRs were not included as they were either just declared (Pugu-Kazimzumbwi and Uzigua) or proposed (Essimingor, Hassama Hill, and Nou).

**Table 1:** Online data sources for the raw data used in this paper: biodiversity (species lists for plants, birds, mammals, reptiles, and amphibians per site); management (revenue generation, tourist numbers, management capacity, and forest disturbance); geospatial data (species range maps, protected areas, etc.) used for the spatial analysis.

| Data | Links |
| --- | --- |
| Raw taxonomic list per site | biodiversity data |
| Data on management of the sites | management data |
| Raw geospatial data | spatial data |

Detailed lists of vertebrate species and plant taxa in the reserves were built upon existing lists for the Eastern Arc Mountains [10, 11, 12, 6], and for coastal forests [13]. Unpublished lists of plant taxa in each Nature Reserve, collated by Roy Gereau of the Missouri Botanical Garden in 2020, were augmented with records from the GBIF portal (accessed in July 2020), and data compiled for other sites using GIS overlay analyses. The designation “taxa” will be used throughout the text to mean “terminal taxa” for plants including species, subspecies, and varieties. In comparison, the unit of analysis for vertebrate will be species only. Species distribution data for vertebrates were compiled using existing GIS data: Birds [14], amphibians and mammals [15], and reptiles [16]. These were intersected with NFR shapefiles provided by the Word Database on Protected Areas, maintained by the United Nations Environment Programme - World Conservation Monitoring Centre (UNEP-WCMC). Additional data sources include published and unpublished reports such as forest management plans and biodiversity survey reports, as well as data from unpublished baseline, mid-term, and end-line NFR surveys undertaken as part of a Global Environment Facility (GEF)-funded project *“Enhancing the Forest Nature Reserves Network for Biodiversity Conservation in Tanzania’*. Data on the taxa occurring in each of the 19 NFRs and 3 proposed NFRs were categorised as endemic (both endemic to Tanzania or to the NFR itself) and threatened with extinction according to IUCN Red List categories: Critically Endangered (CR), Endangered (EN), and Vulnerable (VU). As each taxonomic group had a different total number of taxa, analyses of the proportion of each group in a NFR was carried out separately for each taxonomic category.

Data on the declaration date and area for each NFR were gathered from the Tanzanian government. NFR management effectiveness data came from the Management Effectiveness Tracking Tool (METT) developed by the World Commission on Protected Areas [17]. METT assessments were applied four times, first at the start of the GEF project in 2015; then at project mid-point in 2017; and finally at project endpoint in 2019. A recent assessment of 17 NFR sites was also made in 2021, using the newest methodology METT 4.0. METT scores are interpreted as the higher the score, the better the NFR is managed [18, 19]. Data on income generated and tourism trends were collected from the managers of each NFR, reserve management plans, and Project Implementation Reports (PIR) to the GEF through the UNDP Tanzania country office. In 2019, compiled data were checked by managers of all 17 NFRs and the staff of UNDP-Tanzania and the Tanzania Forest Service [20]. To assess management effectiveness further, data on forest disturbance were collected for 4 NFRs (Amani, Kilombero, Mkingu, and Uluguru) where baseline (2001 - pre NFR creation) and more recent (2019) data exist. Data on disturbance were gathered from 5m on either side of transects within the forest that measured 600 m length (with a total length surveyed of 3.3km; [21]. Data on live, naturally dead, or cut stems of trees were collected. The aim of repeating disturbance transects was to determine whether the declaration of NFR status has had an impact on levels of habitat degradation.

### Data analysis

We assessed the biodiversity value of each reserve by totalling the number of both endemic and threatened taxa known to each reserve. Reserves were then ranked in chronological order of declaration to visualise the change in biodiversity value of the network over time. The correlation between the number of taxa and the date of declaration was tested using the non-parametric Spearman’s rank correlation coefficient as our data were not normally distributed.

The Management Effectiveness Tracking Tool (METT; [17]) was used to assess and compare the effectiveness of each NFR, alongside tourist numbers, income generated, management capacity, and forest disturbance. As previously, our data were not normally distributed; therefore, the possible correlation was tested using the Spearman correlation test in R.

To determine the change in endemic vertebrate species coverage over time with the growing NFR network, we calculated the percentage of endemic species ranges inside and outside the reserves and determined the presence of gap(s) or poorly covered species ranges. Species distribution data from the IUCN Red List for amphibians, reptiles, birds, and mammals were combined with NFR location data to establish species coverage by the NFR network. We grouped NFRs according to the date they were established, resulting in eight groups/time frames (as several reserves were declared the same year). We chose to focus only on endemic species of Tanzania. For each endemic species, the percentage of its range covered by the sites was determined and the results were classified into different categories: (1) 0% of an endemic species’ range covered by the sites: corresponding to a gap species, (2) between 0-2% of an endemic species’ range covered by the sites: corresponding to a poorly covered species, (3) between 2-5%of an endemic species’ range covered by the sites, (4) between 5-10% of an endemic species’ range covered by the sites, (5) between 10-20%, (6) between 20-50%, and (7) more than 50% of an endemic species’ range covered by the sites. The proportion of endemic species was calculated in each of the categories.

We also created species richness maps to visualise the gap analysis. Only the ranges of ‘endemic gap’ species and ‘poorly covered endemic’ species were used. We used the World Database on Protected Areas (accessed in April 2022) to perform a further gap analysis to determine if there are other types of protected areas, in addition to NFRs, that overlap with the gap/poorly covered species still present once all 22 NFRs were included in the analysis. We combined all types of protected areas occurring in Tanzania such as National Parks, Nature Reserves, Game Reserves, Forest Reserves, Conservation Areas, Village Land Forest Reserves, and Game Controlled Areas. We determined the percentage of species range covered by the different types of protected areas and computed the proportion of species inside the classified categories. Analyses were carried out using the statistical software R (version 4.0.5) [22], the spatial Geographic Information System QGIS (version 3.16.14), and Microsoft Excel.

## Results

### Development of the NFR network over time and coverage of Tanzanian biodiversity values

We found that older NFR sites support a larger number of recorded threatened taxa when compared to recently declared sites (Figure 2A and Table 3). Amani, Kilombero, and Uluguru NFRs have the highest number of threatened taxa, followed by Nilo, Chome, Mkingu, Uzungwa Scarp, Rondo, and Magamba. Many of these species are also endemic to Tanzania (see Figure 2B, Amani to Mangula). Predictably, as new reserves were added to the network there was a gradual increase in the number taxa in the NFR network, but this was found to plateau for all reserves declared since 2019 (Figure 2C). A significant positive correlation was found between the cumulative numbers of endemic taxa, total taxa, and date of declaration of the reserves (Figure 2, Spearman correlation test, rho = 0.97, p <0.001 for all taxon groups). This overall significant increase is also found in each taxon group separately (Spearman correlation test, rho = 0.95, p < 0.001 for each taxon group). We found no significant correlation between the year of establishment and the reserve area (p = 0.8755).

**Fig 2:**
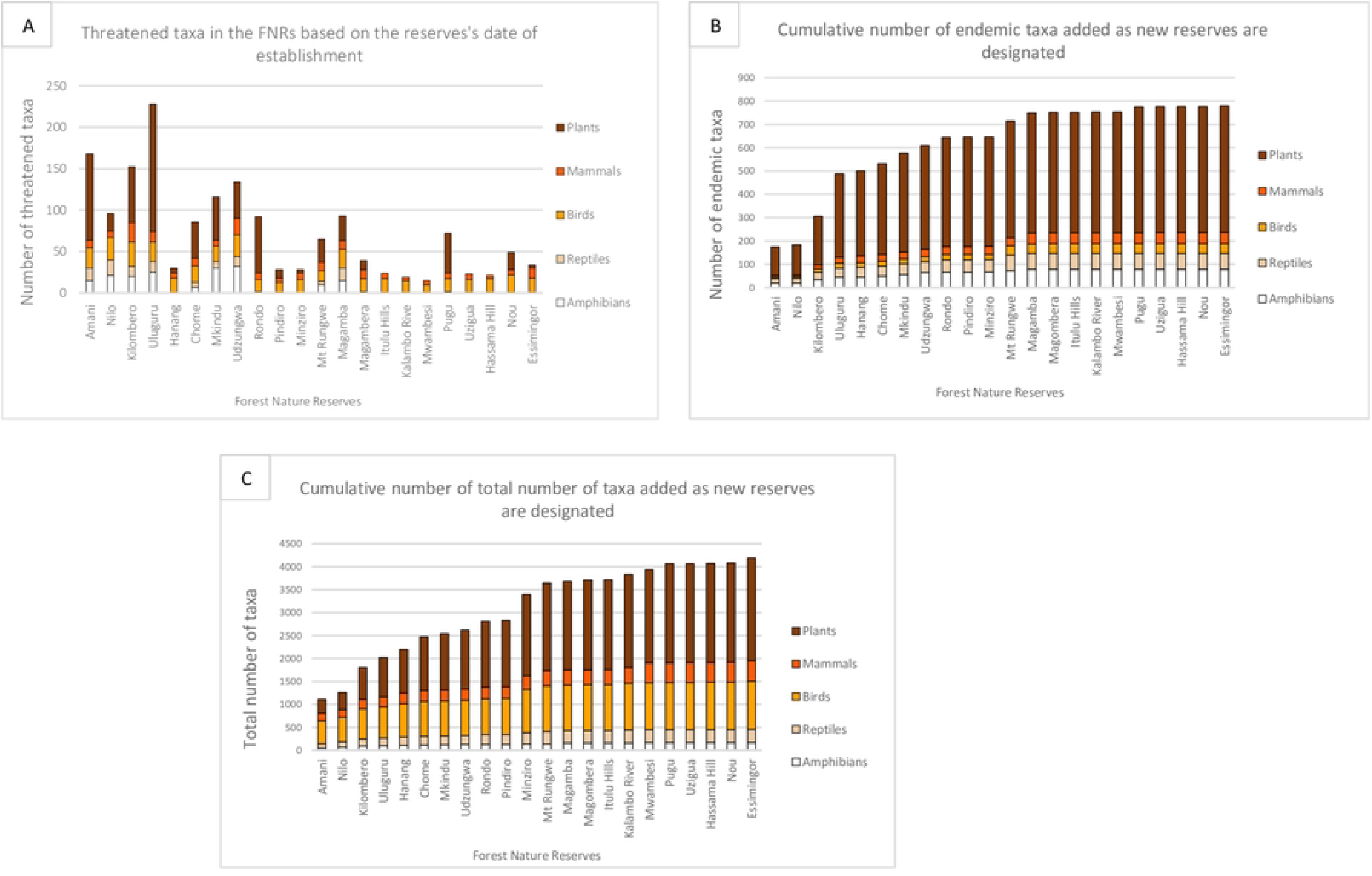
Importance of the network of NFRs in terms of Tanzanian endemic and globally threatened plant and vertebrate taxa as the network has been expanded since 1977. A: Number of threatened taxa in the NFRs. B: Cumulative number of endemic taxa. C: Cumulative number of total taxa (including endemic, threatened, and other taxa). Reserves are ordered according to the year of their designation (the last three reserves are proposed (no year of designation)).

### Effectiveness of NFR management in achieving conservation goals

Mean METT scores for the NFR network increased over time. The overall mean percentage METT score for the 11 NFRs present in 2015 was 55%, 69% in 2017 for 12 NFRs, 69% in 2019 for 17 NFRs assessed, and increasing in 2021 to a mean score of 86.8% for the 17 NFRs evaluated (Figure 3A). However, the Spearman correlation test did not find any significant relationship between the date and the average final METT scores across the reserves (Figure 3A). NFRs with high numbers of endemic taxa were found to have higher management effectiveness (Figure 3B), but the relationship was not significant (Spearman correlation test, effect size=0.03, p = 0.46) (Figure 3B). The total tourist numbers (mainly international) steadily increased in most NFRs, from 1698 tourists in 2016 to 4097 tourists in 2019 (Figure 3C). The positive correlation between years and visitor numbers was significant (Spearman correlation test, effect size= 0.95, all p values < 0.0123, N = 17) (Figure 3: C). However, income generated (US dollars fixed across the four years measured) decreased slightly from 2017 to 2019 (Figure 3: D). Furthermore, no significant relationship was found between the average income generated and the four years studied, except between 2017 and 2021 (Spearman correlation test, effect size= 0.8095, p value = 0.02178, N = 17) (Figure 3: D). Management capacity including the number of staff, buildings, transport, and equipment’s present in each NFR increased between 2015 and 2019 except staffing (number of staff and rangers), which decreased in 2019 (Figure 3: E). Only large equipment (corresponding to the number of computers, photocopier scanner, printers, GPS units, solar batteries, and hard drives) displayed a significant increase between 2015 and 2019 (Spearman correlation test, rho = 0.61, p value = 0.03669, N = 17).

**Fig 3:**
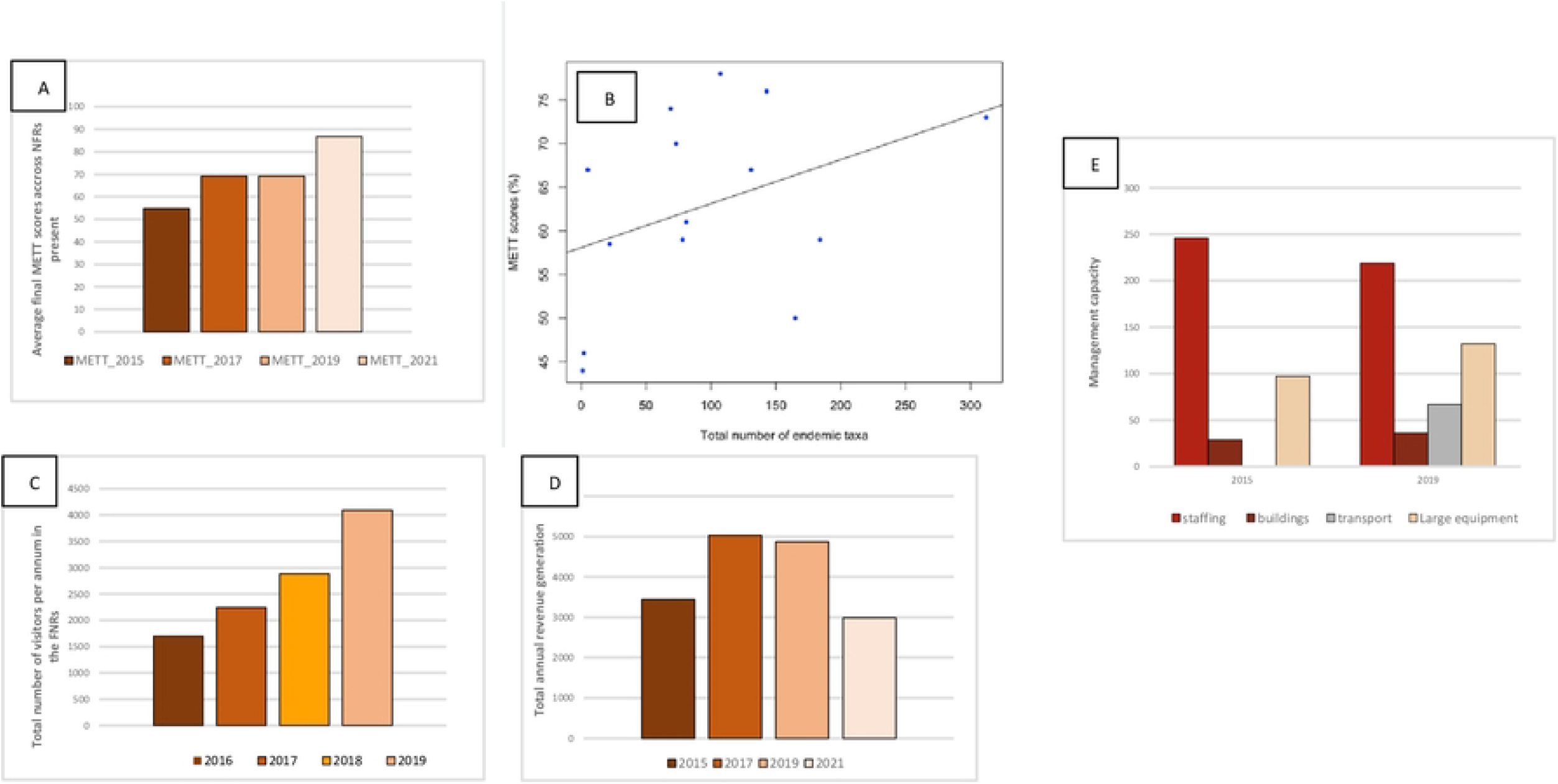
Changes in management data from 2015-2021 for: A) average final percentage METT score across the sites; B) relationship between average METT score and biodiversity value ranked in ascending order (linear trend line not significant); C) annual number of tourists in all sites (Table S1 for full data); D) average total revenue generated in the reserves (US dollars); E) sum of the management capacity in 2015 and 2019 comprising staffing (staff and rangers), buildings (offices and ranger posts), transport (vehicles), and large equipment.

Analysis of forest disturbance transect data from four NFRs in 2001 (before these NFRs were declared - except for Amani) and 2019 shows that numbers of cut trees and cut poles per hectare have declined since the reserves were declared (except for Uluguru and Kilombero NFRs) (Figure 4). This suggests that improved management at these sites has prevented some harvesting of woody products from the NFRs. The number of live trees was stable (in Uluguru NFR) or decreased (in Kilombero and Mkingu NFRs), while the number of live poles increased (expect for Kilombero NFR), suggesting that illegal logging has continued in some areas and that this may have promoted regeneration of smaller trees and bushes in the various sites.

**Fig 4:**
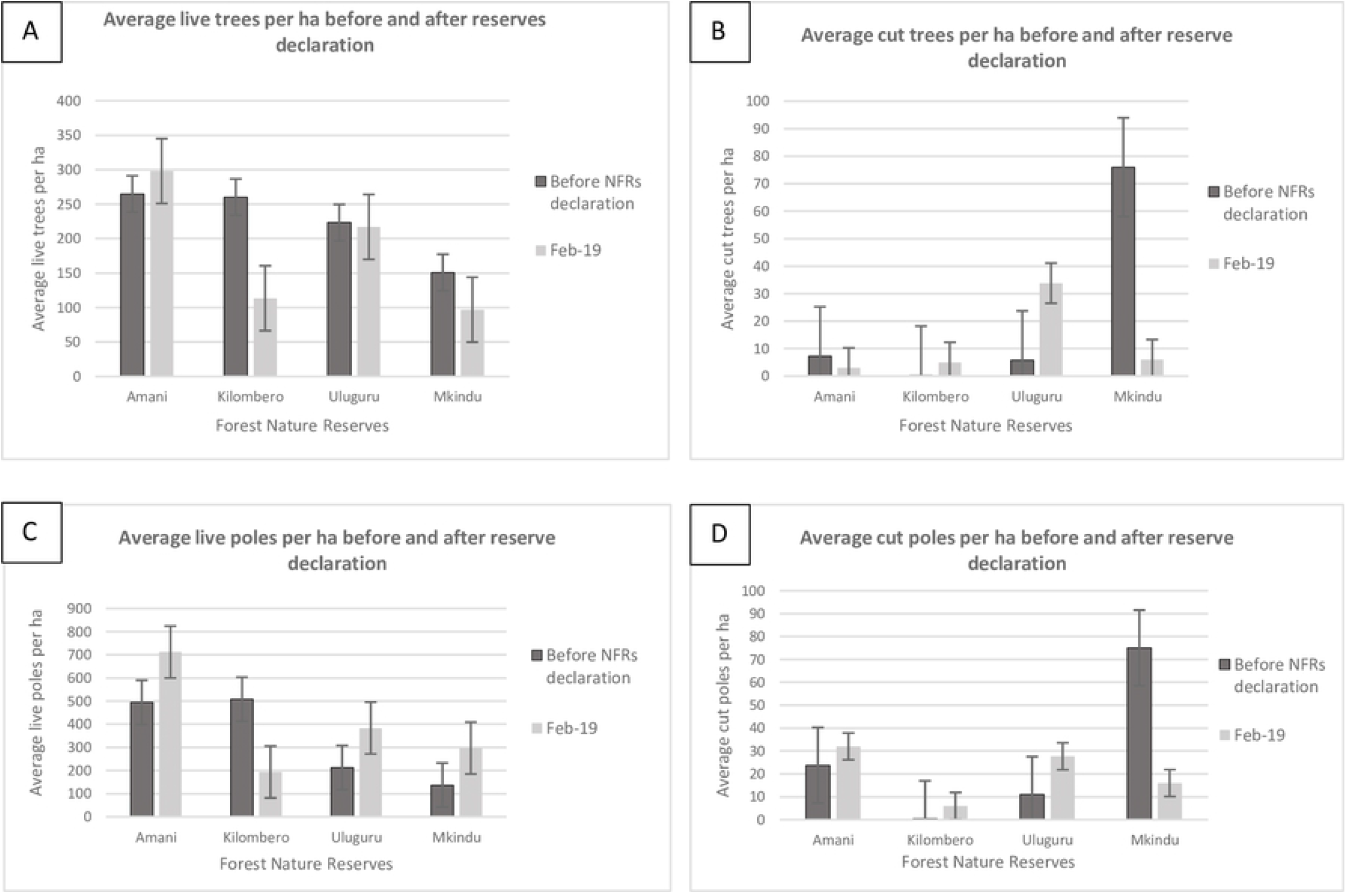
Changes in forest extraction activities in terms of cutting of: A) living trees (>20 cm dbh), B) cut trees, C) living poles (<20 cm dbh), and D) cut poles in Amani, Kilombero, Uluguru, and Mkingu Nature Forest Reserves before reserve declaration and in February 2019.

### Coverage of endemic and threatened species by the NFR network

GIS analyses show that the addition of sites over time decreased the number of Tanzanian endemic vertebrate species missing from the reserve network (gap species). For Tanzanian endemic bird species, the gap species were covered after the addition of the last sites of the network (Figure 5: A). However, the proportion of poorly covered endemic bird species (between 0-2% of their range covered) was high and was the highest across all of the endemic species. For species with a larger range coverage such as mammals, the gap declined rapidly to reach 5% when all sites were included in the analysis (Figure 5: B). When the full network was considered, the proportion of endemic amphibian species with more than 50% of their range protected was higher than other taxa, with more than 25% of endemic amphibian species ranges partially covered by the NFRs (Figure 5: C). Nevertheless, amphibians had the highest proportion of gap species considering all the sites (Figure 5: C). As for the other vertebrate groups, the proportion of endemic reptile gap species is decreasing, as new sites were added to the network to reach less than 10% of gap species considering all the NFRs (Figure 5: D).

**Fig 5:**
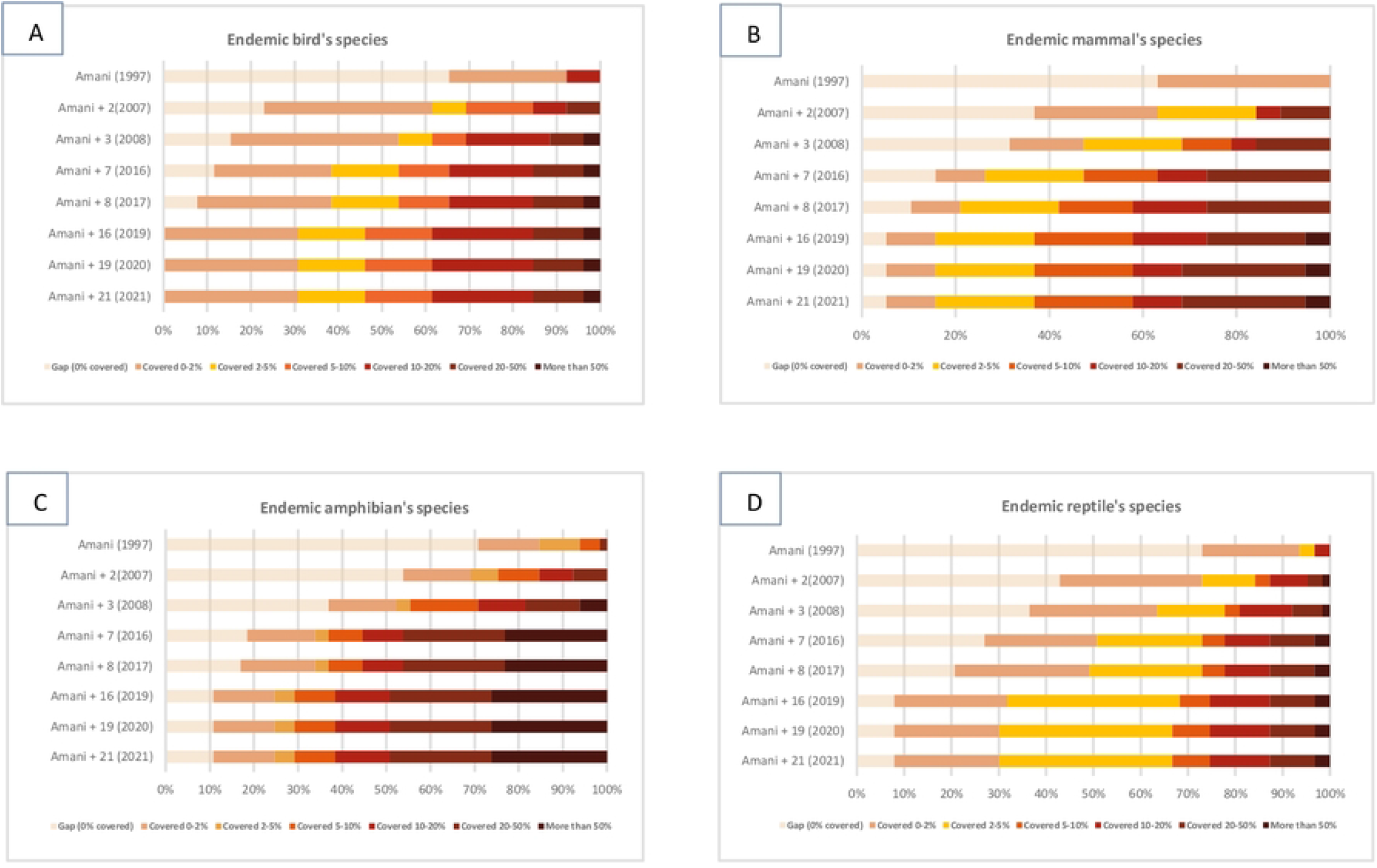
Proportion of Tanzanian endemic vertebrate species ranges covered by NFRs over time. Years when the reserves were gazetted are in parentheses and the number of reserves declared in each time frame is indicated after “Amani”. The light beige bars refer to the endemic gap species, where 0 percent of their range is covered. Other colours show the percentage of endemic species range covered by reserves. Scale is from light brown, where between 0 and 2 percent is covered, yellow, between 2 and 5 percent covered, orange, between 5 and 10 percent, dark red, between 10 and 20 percent, brown, between 20 and 50 percent, and finally dark brown, where more than 50 percent of the species range is covered by the reserve.

Richness maps of gap species endemic to Tanzania (including all Tanzania endemic species in the four taxonomic groups of vertebrates studied) show, not surprisingly, a decline in the proportion of gap species as more reserves were added to the network. However, there is a lack of coverage within the NFR network for endemic reptile gap species in the South-East part of the country around Rondo and Pindiro NFRs, and in some central areas of the country.

Including all types of protected areas present in Tanzania, the proportion of gap and poorly covered species decreased strongly in comparison to only NFRs (Figure 7). No poorly covered species were found considering all the protected areas, compared to 30% of poorly covered bird species considering only the NFRs (Table 2). Almost 7% of endemic amphibian species remain unprotected, compared with approximately 11 % in NFRs (Table 2).

**Table 2:**
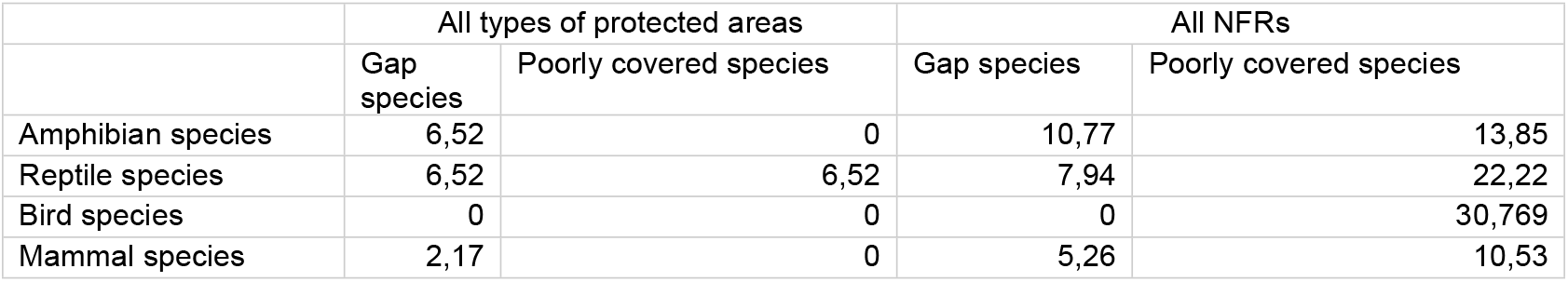
Comparison of the proportion of gap and poorly covered endemic amphibian, reptile, bird, and mammal species, when all protected areas are included compared to only the NFRs

|  | All types of protected areas |  | All NFRs |  |
| --- | --- | --- | --- | --- |
|  | Gap species | Poorly covered species | Gap species | Poorly covered species |
| Amphibian species | 6,52 | 0 | 10,77 | 13,85 |
| Reptile species | 6,52 | 6,52 | 7,94 | 22,22 |
| Bird species | 0 | 0 | 0 | 30,769 |
| Mammal species | 2,17 | 0 | 5,26 | 10,53 |

**Table 3:**
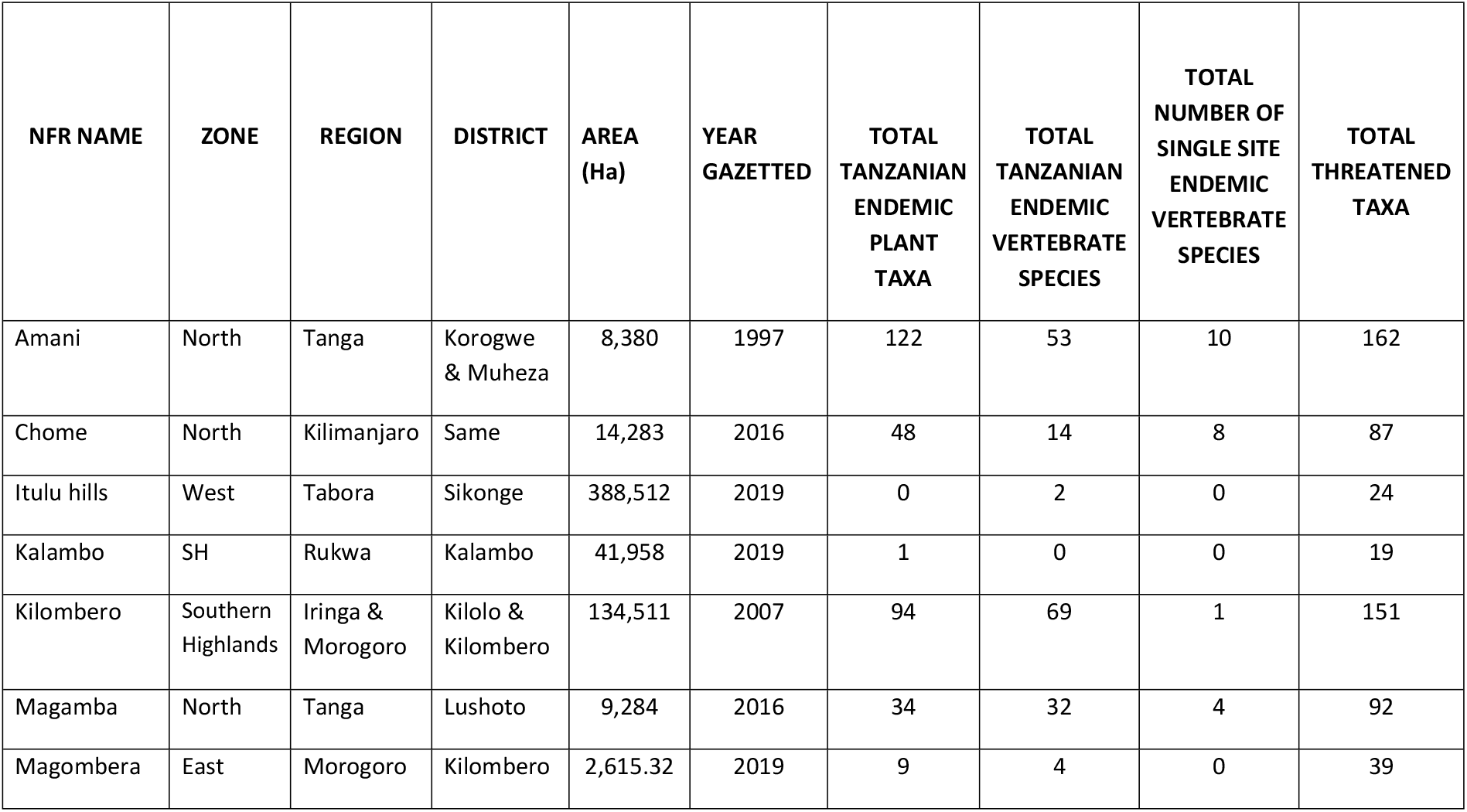
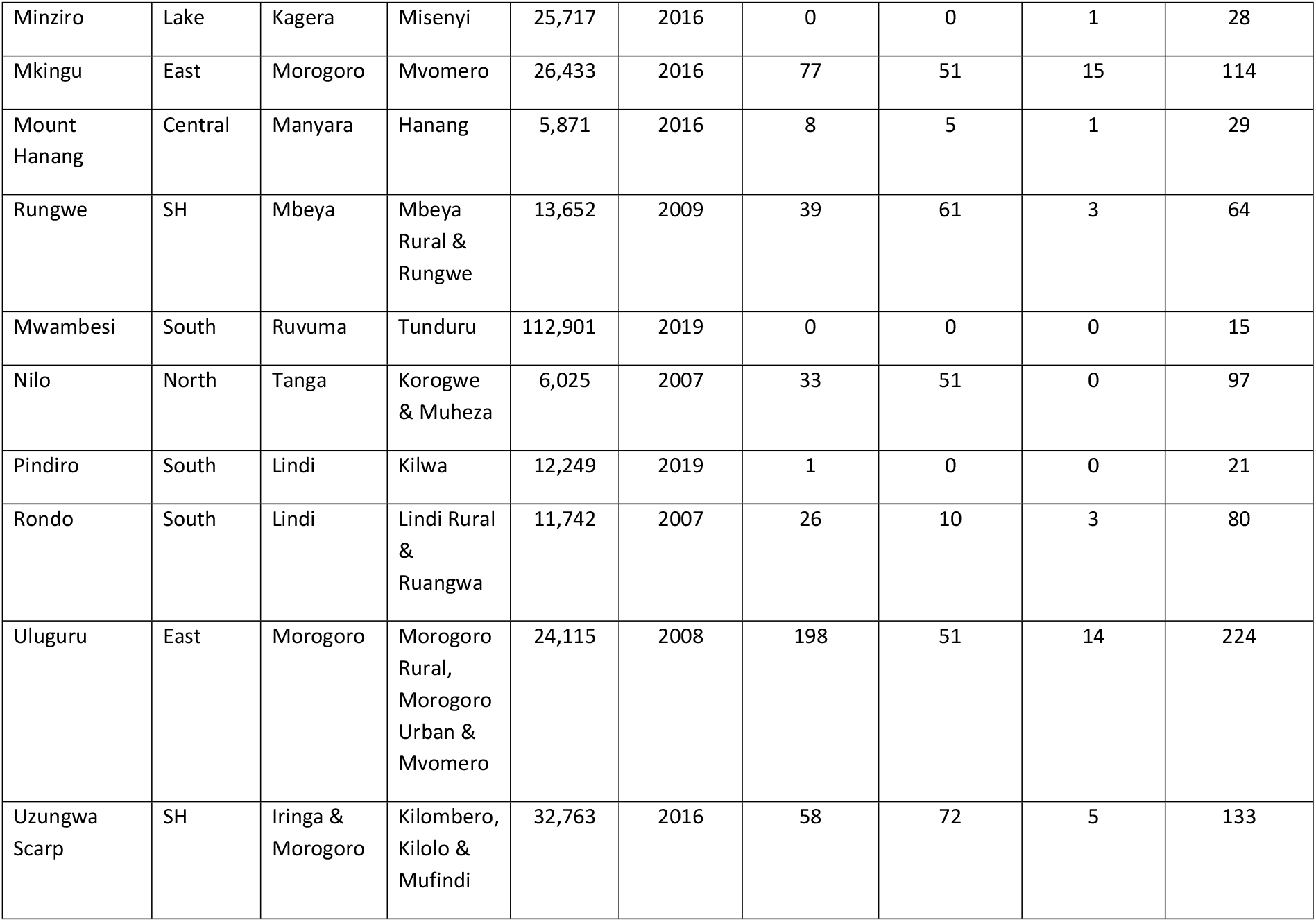
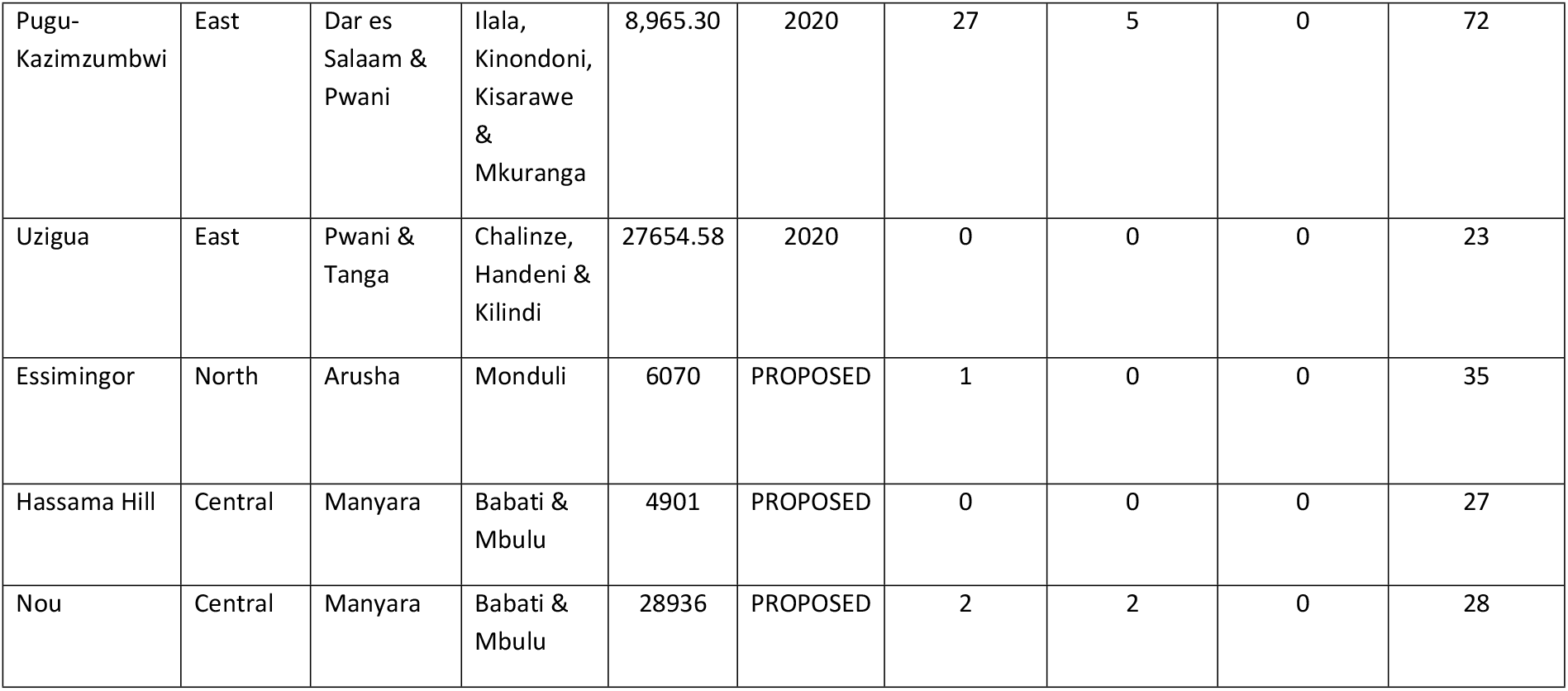
Basic attributes of 19 declared and three proposed Tanzanian Forest Nature Reserves

## Discussion

### How well do NFRs capture Tanzanian biodiversity values?

The NFR network was not formally designed using conservation planning approaches, but using a combination of expert knowledge, the framework of ecoregions, and picking the ‘best’ remaining sites owned by the current management body – Tanzania Forest Service (formerly Forestry and Beekeeping Division of the Ministry of Natural Resources and Tourism). The first designated NFR sites, found in the Eastern Arc Mountains or Coastal Forests ecoregions, have higher biodiversity values than more recently declared sites. These regions are known to have globally important biodiversity values [23, 24, 10], but have also received higher research funding and survey efforts [25, 10] and were the first to be protected, with other sites in other ecoregions following as resources/data became available. Many of the newer NFRs have received little or no biodiversity survey attention, so we expect their known values to increase if they receive similar biodiversity survey attention as the older NFRs (see e.g., Minziro NFR, [26]). Nevertheless, the recently designated sites also contain valuable biodiversity, and contribute to an increase of the biodiversity values within the reserve network over time (Figures 2: B, C, D). The network of reserves now covers most of the endemic and threatened vertebrate species in Tanzania, however, not their full ranges. The most important sites in terms of biodiversity value also show slightly higher METT scores, suggesting that resources have been allocated to more important sites (Figure 3: B), even though the increase is small.

### Do gaps remain in the NFR network?

As more reserves have been added to the NFR network over time, the proportion of gap species has decreased for all species groups (Figure 5). This is especially true for endemic mammal and bird species, which respectively had 5% and 0 % of gap species remaining after all NFRs were added to the network. However, the proportion of poorly covered species increased for some groups such as birds, which had more than 30% of species poorly covered including in all the NFRs compared to 26% including only Amani NFR (Figure 5). Due to the relatively small size of the NFRs and the large range of some Tanzanian endemic species, NFRs could not adequately cover the ranges of all the endemic species in the country. To achieve a larger coverage of the ranges of these species we needed to include other types of terrestrial protected areas in our analysis (Figure 7) [27].

The major NFR gap area identified was in the coastal forests (Figures 6 and 7). Even after adding all the types of protected areas present in the country, the gap in the South-East part of the country is still present. There are few and only relatively small forest reserves present in that area. Further assessment is needed to determine the need for additional reserves to cover distributions of the gap species. Perhaps one of the small forest reserves present in the area (such as Rondo or Matapwa Forest Reserve) could be expanded to cover a part of the gap species’ ranges, or a new type of reserve could be implemented.

**Fig 6:**
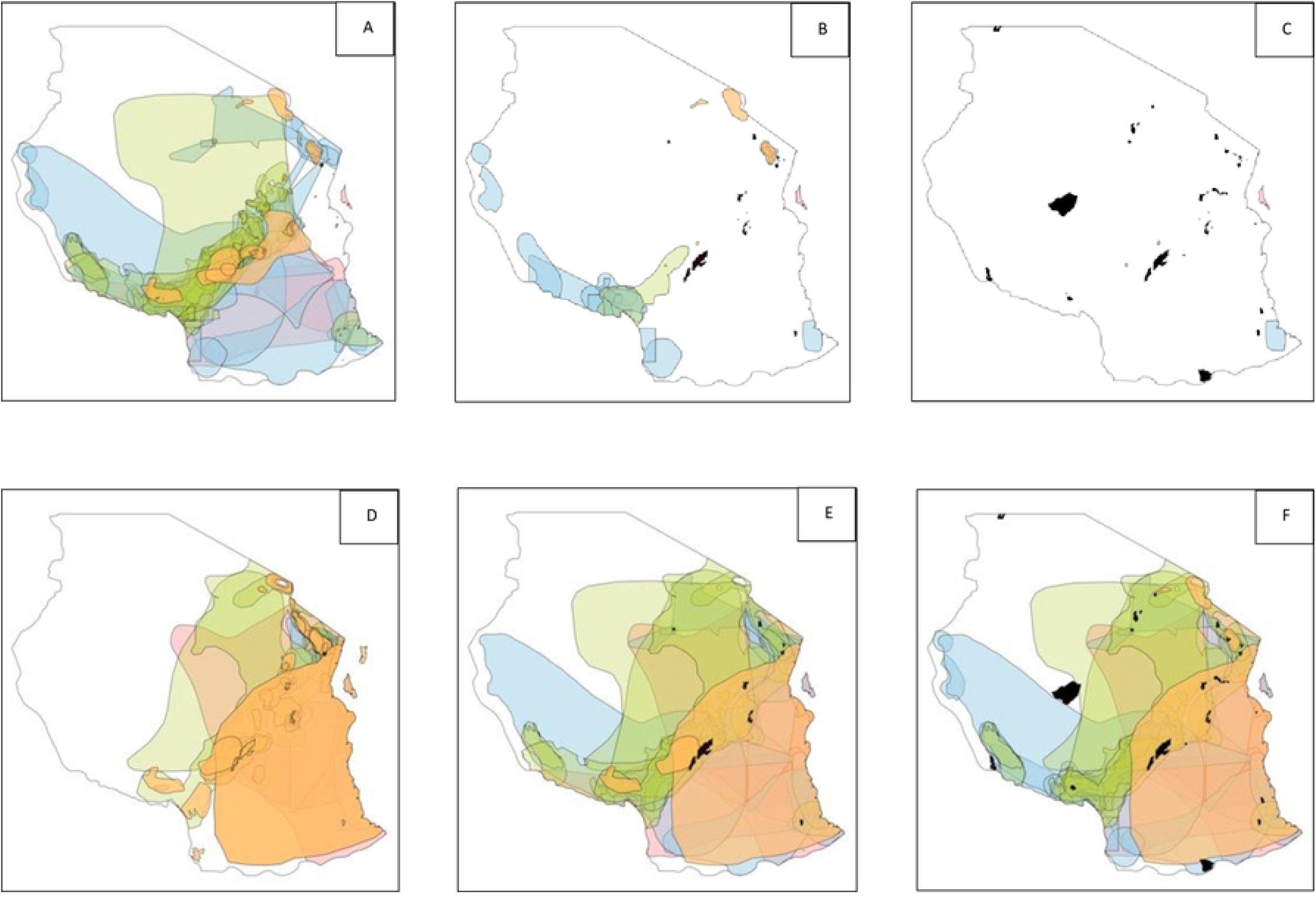
Coverage of the range of Tanzanian endemic ‘gap’ species (panels A, B, C) and poorly covered endemic species (panels D, E, F). Panels A and D show endemic species coverage in 1997, when the first reserve was added to the network. Panels B and E show the endemic species coverage in 2017, with 9 reserves in the network. Panels C and F represent the species coverage in 2022, with 19 NFRs and 3 proposed NFRs included in the network. The orange coverage range displays endemic mammal species, blue reptile species, green bird species, and light pink amphibian species. The NFRs are indicated as black polygons.

**Fig 7:**
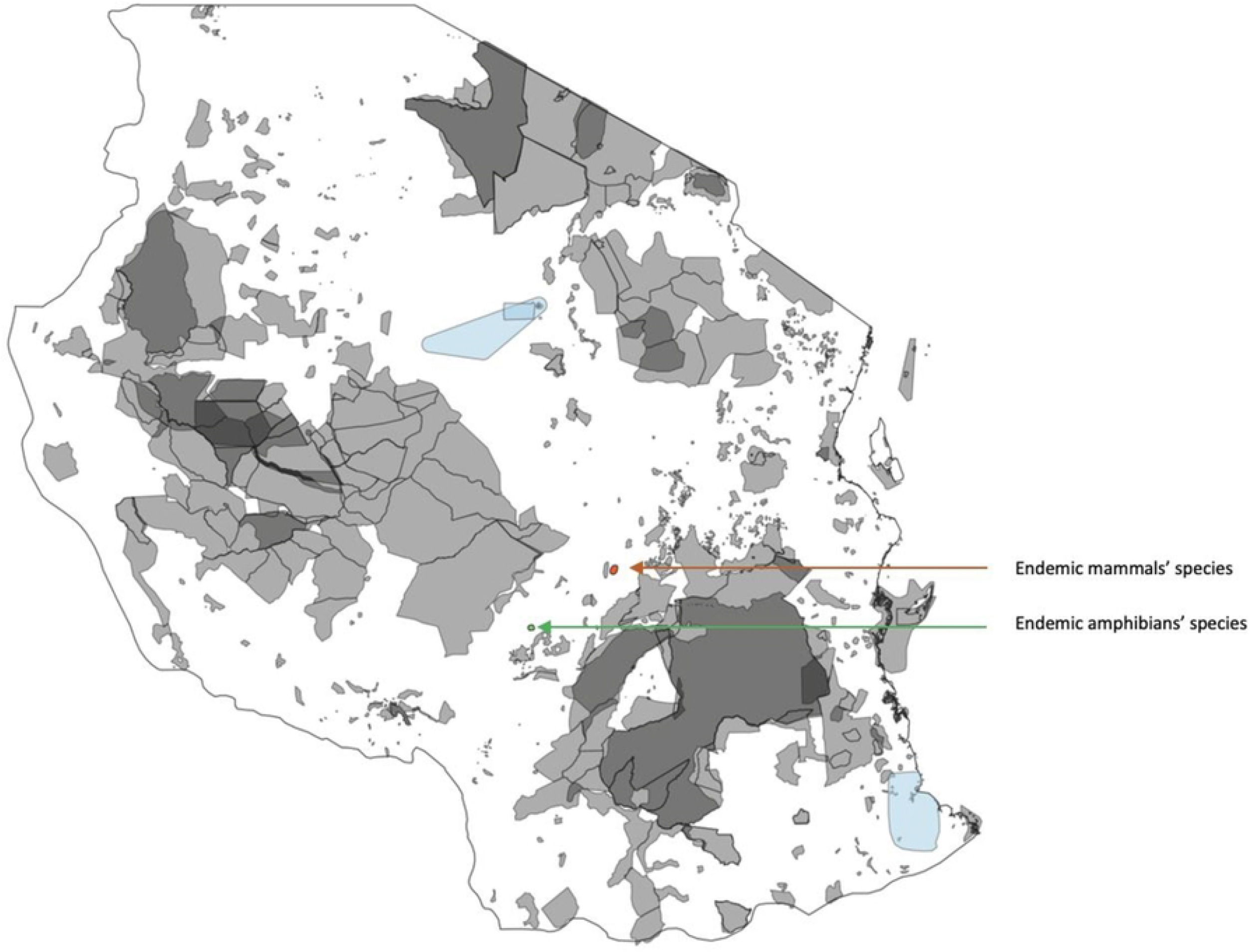
Coverage of the ranges of gap and poorly covered endemic Tanzanian vertebrate species by the protected area network in Tanzania. Orange shows endemic mammal species, blue reptiles, and green represents endemic amphibian species. There are no gap species of birds. Protected areas are indicated as grey polygons.

### Management of the Nature Forest Reserves

The NFR network in Tanzania has expanded over the past 25 years from a single site to 19 sites covering the main forest types in the country. This represents an invaluable asset, especially considering that the world-renowned network of National Parks in Tanzania mainly protects non-forest or drier forest habitats, with a few notable exceptions that include Udzungwa Mountains, Mahale, and Kilimanjaro NPs. The future of the NFR network depends on the policies and decisions of the Tanzanian government, but also on the ways in which the sites are funded and managed. Development of the network has been funded by a combination of the Tanzanian government (TFS) and donor funding including the GEF, the Eastern Arc Mountains Conservation Endowment Fund, and private donors such as the Aage V. Jensen Foundation, as well as local and international NGOs. Thanks in part to these investments, there is evidence of management effectiveness improvement (METT scores) from 2015 to 2021 (Figure 3: A).

The number of tourists in the sites increased from 2015 to 2019, but the contribution to management costs remains small (Figure 3: C). Tourism has primarily been by visitors from foreign countries, yet the number of Tanzanian tourists has recently increased, especially during the COVID-19 pandemic. Older reserves such as Chome, Magamba, and Amani, with better infrastructure, accessibility, and proximity to the northern tourism circuit, had the largest growth in tourism numbers. However, reported direct income from tourism in the NFRs remained low, ranging from USD 3447 in 2015 to USD 5032 in 2017 to less than USD 5000 in 2019 (Figure 3: D). Since 2020, international tourism has been significantly impacted by the COVID-19 pandemic, which financially undermines these reserves as well as other protected areas in the country. There remains potential for improving local tourism visitation to these NFRs and increasing revenue collection, but the potential to pay the full management costs remains low and care must be taken not to develop facilities that remain under used.

The fact that forest disturbance (using the proxy of cut trees and poles) generally declined from 2001 to 2019 in three of the four studied NFRs is suggestive of improved management at these sites. However, this assessment is only available for four reserves, and we have no overview of how this condition may have changed in other sites. Anecdotal evidence suggests that some other reserves are suffering considerable disturbance, including hunting pressure, that is not captured in the data we have available. This is the case in Uzungwa Scarp NFR, where decades of poor management of the former Forest Reserve have led to dramatic loss of wildlife [28, 29]. However, since its upgrading to NFR status in 2016, increased management efforts supported by local and international agencies and implemented in close partnership with TFS have led to a considerable improvement [30]. Key strategies have included capacity building of patrol staff through training workshops and provision of patrol gear, engagement of local community members in collaborative forest patrols, use of novel technologies (such as smartphones to collect patrol data and camera traps for surveillance), and regulated access by communities to non-timber forest products such as firewood, medicinal plants, mushrooms, worshipping sites, and the use of trails for traveling across areas. A similar process has recently begun to show positive results in the Kilombero NFR. Hunting pressure and long-term forest encroachment may have wiped out the endemic Udzungwa forest partridge species from the Nyumbanitu forest within the Kilombero NFR, where it was present in 2016 [31]. This was one of the only two forest fragments globally where the species occurred, both in the Kilombero NFR. However, law enforcement and awareness raising in local villages have started since August 2021, which may slowly start improving the conservation status of this and other single-site endemic species.

### What management and research suggestions can we offer?

To improve the quality of management of NFRs, we suggest that future management plans should place an emphasis on enforcing hunting and logging regulations in all sites, raising awareness in villages surrounding reserves and providing local people with viable alternatives to activities that impact the forests and their biodiversity. The example of schemes such as those mentioned for the reserves in the Udzungwa Mountains in which TFS has signed agreements with local and international NGOs to support capacity building and facilitate management, including the engagement of local community members in intelligence and forest patrol efforts, may represent a good model forward. Furthermore, we recommend that future management plans include an expansion of the current network, incorporating the endemic gap species and poorly covered species as well as other unprotected forest taxa still present in the country. Management planning should focus on forest extension, creation, and implementation of meaningful Joint Forest Management regimes between TFS and surrounding communities, improvement of forest quality, and more sustainable use of its resources.

Biodiversity and management effectiveness monitoring of current and future sites will be important to continue to understand the coverage and efficacy of Tanzania’s NFR network. For example, an equal number of surveys for all taxa and protected areas is proposed to be accomplished, beginning with NFRs with little or no biodiversity data (e.g., Hassama, Itulu Hill, Mwambesi, Pindiro, and Uzigua NFRs, all virtually unknown in terms of plant data). We recommend that additional surveys on species/taxon distribution patterns be performed, and that future research include analyses of taxon distribution fluctuation due to global warming [32]. Furthermore, life history, human population growth, and species abundance data should be added to future analyses [33].

The implementation of a stable budget through the years to afford new equipment, facilities, and infrastructure, and to increase the patrols in the NFR network is recommended. To increase the budget available to a reserve, we suggest the following: capturing revenue from ecosystem services (such as water catchment and carbon stocking), soliciting additional investment plans or donations, and increasing the revenue generated from touristic activities [10]. Furthermore, we recommend that the TFS management consider a further expansion of the current network to include the (limited number of) endemic gap and poorly covered species that are outside the NFR network but could be included with some management status changes for existing sites.

## Limitations

The data and analyses used here are subject to several limitations. Firstly, it was difficult to assure the quality of METT scores, except METT Version.4.0, and data on recently declared NFRs were missing. This reflects earlier studies in which METT data have been valuable but need to be interpreted with care when done using self-assessment approaches [17, 34], especially for older METT versions, i.e., METT 1-3. To obtain the best outcome from the METT assessment process, studies suggest the application of guidelines for the METT assessment and involvement of a group of people in the assessment to ensure that results are unbiased [19, 35). This is also more required by METT 4; such that local communities must be involved in the assessment. Second, only spatial data on the range of species were used, without life history or species population data being available. This means that we cannot assess the viability of populations within a NFR or within our broader gap analysis. Species range data may also be biased, especially in areas situated farther from protected areas, which may have been poorly sampled [27]. Furthermore, defining areas of current species occurrence may be misleading and might lead to ineffective protection due to the discontinuous distribution of species due to climate change [32]. Third, ecological data on current key local threats impacting nature reserves and the taxa were not available in a standardized fashion across the network, with only partially complete data available up to 2019. The COVID-19 pandemic will also have impacted the management and functioning of NFRs in Tanzania, but updated data are not available.

## Acknowledgements

We thank the many staff of the Tanzania Forest Service and key donors, including the Global Environment Facility (via UNDP and the World Bank), the Critical Ecosystem Partnership Fund, European Union, Governments of Norway, Finland and Denmark, and facilitation by several NGOs: WWF, Tanzania Forest Conservation Group, CARE International, IUCN, WCS, BirdLife International, and African Wildlife Foundation.

